# Catalytically inactive SHP1-C453S mutant gain of “robust LLPS” function

**DOI:** 10.1101/2024.03.06.583685

**Authors:** Qichen Zhang, Tianyue Sun, Qi Chen, Guangya Zhu, Xiangyu Kong, Yiqi Du

## Abstract

SHP1 is a non-receptor protein tyrosine phosphatase extensively expressed in hematopoietic cells, exerting a pivotal role as an immunosuppressive factor. Our previous studies have suggested that SHP1 can undergo liquid-liquid phase separation (LLPS). In this study, the SHP1-C455S mutant, commonly utilized in biochemical assays due to its lack of catalytic phosphatase activity, unexpectedly exhibited a remarkably robust ability for LLPS. Since the C453S mutation has been previously shown to potentially induce a conformational transition of SHP1 from a closed to an open state, we hypothesize that the enhanced LLPS capability of SHP1 may be facilitated by this conformational alteration. The SHP1-C453S mutant exhibited robust LLPS activity, while completely abrogating its phosphatase activity. This allows for effective investigation of the catalytic activity and LLPS capability of SHP1.

## Introduction

SHP1, encoded by the PTPN6 gene, is a non-receptor tyrosine phosphatase predominantly expressed in hematopoietic lineages including leukocytes, neutrophils, and immune cells, while it is expressed at low levels in epithelial and endothelial cells^[1,2]^. SHP1 is reported to reduce the duration and amplitude of downstream signaling molecule cascades following immuno-receptor activation^[3,4]^. Moreover, SHP1 is essential to immune cell development, function and differentiation and has been established as a critical immune checkpoint and a potential target for therapeutic intervention^[5]^.

The concept of liquid-liquid phase separation (LLPS), a physicochemical phenomenon that results in the segregation of binary or multicomponent mixtures into distinct phases under specific conditions, has recently gained significant attention within the field of biological science. LLPS is now recognized as a fundamental mechanism governing numerous biological processes and facilitating the formation of biomolecular condensates, which generate organized structures capable of concentrating and compartmentalizing intracellular biochemical reactions^[6,7]^. The phase-separation ability of SHP1 has been demonstrated in our previous studies^[8]^. In this study, we investigated the LLPS capacity of SHP1-C453S, a commonly utilized catalytically inactive mutant. Our findings demonstrate that the C453S mutation in SHP1 confers exceptional LLPS capability, surpassing even that of the previously studied strong phase-separating protein SHP1-E76A.

## Materials and methods

### 2.1. Protein expression and purification

The full-length WT SHP1 gene was obtained from a human cDNA library and was amplified using PCR. Standard PCR was used to construct a series of SHP1 mutants, which were confirmed by DNA sequencing. The pET28a vector was used to insert all constructs of human SHP1 6x histidine tag coding sequence was added to the N terminus of the constructs. Escherichia coli BL21 (DE3) cells were grown and cultivated in a lysogeny broth medium. Protein expression was induced at OD 0.4-0.6 using 0.5 mM isopropyl-beta-D-thiogalactopyranoside and allowed to grow overnight in the dark at 16°C. After centrifugation at 4,000 rpm for 30 min at 4°C, the pellets were then lysed by sonication on ice in a buffer containing 300 mM NaCl, 100 mM Tris-HCl (pH 8.0), 15 mM imidazole, 0.1% Triton X-100, 1 mM Tris(2-carboxyethyl)phosphine and 1 mM phenylmethylsulfonyl fluoride. The obtained residue was subjected to super centrifugation at 10,000 rpm for 20 min at 4°C. The resultant supernatant was loaded onto a His Trap HP chelating column(Cytiva) and eluted in a buffer containing 300 mM NaCl, 100 mM Tris-HCl (pH 8.0) and 100 mM imidazole. Using a buffer containing 150 mM NaCl and 25 mM HEPES (pH 8.0), the fractions containing SHP1 were concentrated and loaded onto a HiLoad 16/600 Superdex 200 PG column(Cytiva). Proteins were identified by sodium dodecyl sulfate polyacrylamide gel electrophoresis, the harvested fractions were collected, and the proteins were condensed to 10 mg/mL or more and stored at -80°C.

### 2.2. In vitro LLPS assay

For image shootting, SHP1-WT, SHP1-C455S and SHP1-E76A proteins were diluted to a final estimated concentration of 40 μM in a buffer containing 25 mM HEPES (pH 8.0), 50 mM NaCl with or without 6.7% PEG20000. The mixture was incubated at room temperature for 5 min and 1 μL of each sample was pipetted onto a glass dish and imaged using THUNDER Imaging Systems (Leica Microsystems).

For Turbidity test, SHP1-WT, SHP1-C455S and SHP1-E76A proteins were diluted to a final estimated concentration of 0-8 μM in a buffer containing 25 mM HEPES (pH 8.0), 10-50 mM NaCl with or without 4% PEG20000. The mixture was incubated at room temperature for 30 min and 30 μL of each sample was added onto a 384-well white polystyrene plate with a clear flat bottom. The OD600 nm was measured using Multiskan SkyHigh Microplate Spectrophotometer (Thermo Fisher Scientific).

### 2.3. SHP1 phosphatase assay

SHP1 catalytic phosphatase activity was monitored using p-Nitrophenyl phosphate (pNPP, 333338-18-4, D&B). The reactions were performed in a 384-well white polystyrene plate with a clear, flat bottom (Corning) in a final volume of 30 μL at room temperature. Purified SHP1 proteins from WT and mutant were diluted to 8 μM in a buffer containing 25 mM HEPES (pH 8.0) and 150 mM NaCl. The pNPP substrate was then added into the system to start the enzymatic reaction at a final concentration of 10 mM, and absorbance of the product was measured at OD405 nm every 10 s for 10 min using Multiskan SkyHigh Microplate Spectrophotometer (Thermo Fisher Scientific).

### 2.4. Data analysis

Data were analyzed using Graphpad Prism (version 8.0.2; GraphPad Software, Inc., La Jolla, CA, USA) and presented as means ± Standard Deviation (SD) (n = 3 experiments).

## Results

SHP1 comprises two SH2 domains, one catalytic PTP domain, and a C-terminal tail (Figure 1A). The initial goal of this work was to obtain three proteins: SHP1-WT (wild type), the inactive mutant SHP1-C453S, and the active mutant SHP1-E76A, to serve as controls for subsequent studies on SHP1 catalytic phosphatase activity. In this study, we used the same method to purify these three proteins in E. coli, achieving monomeric protein (protein concentration >90%), with the gel filtration result of C453S serving as a representative image (Figure 1B). During the purification process, we were surprised to find that the SHP1-C453S mutant became turbid at room temperature (Figure 1C-D), a state similar to that of SHP1-WT in PEG buffer, suggesting that the SHP1-C453S mutant might have a strong capability for phase separation. To confirm the phenotype of phase separation, we directly observed the LLPS of SHP1 by a microscope. At the protein concentration of 40μM, SHP1-C453S formed noticeable round droplets and underwent fusion, displaying classic LLPS characteristics, whereas almost no droplets were observed for the WT (Figure 2A). To further quantify the differences in LLPS between the mutants and WT, turbidity was measured at different protein and salt concentrations by spectrophotometer. The bivariate plots analysis of in vitro LLPS assay showed that SHP1-C453S indeed exhibited robust LLPS activity, which was stronger than SHP1-E76A (Figure 2A). LLPS assay with PEG also showed a similar phenomenon (Figure 2B). Finally, phosphatase catalytic activity was tested using the pNPP as substrate, the results of enzymatic assays revealed that SHP1-E76A mutation resulted in a significant increase in phosphatase catalytic activity, while SHP1-C453S led to a complete loss of enzyme activity, consistent with previous reports (Figure 2C).

**Figure 1.**
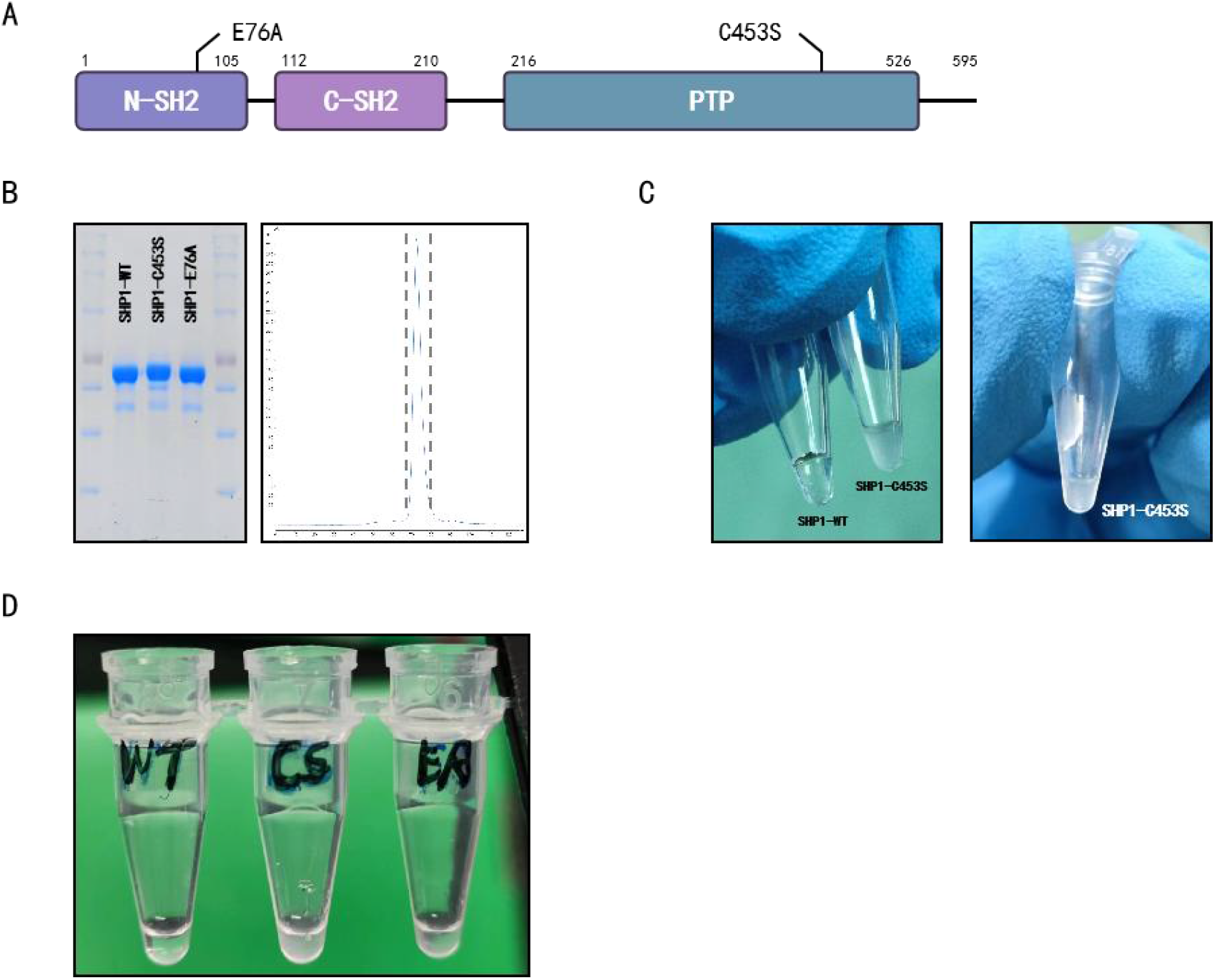
Purification and Turbidity Measurement of Three Proteins: SHP1-WT, SHP1-E76A, and SHP1-C453S. 1A: Schematic diagram of SHP1 protein structure and mutation sites. 1B: Gel electrophoresis images of proteins (SHP1-WT, SHP1-C453S, and SHP1-E76A) and gel filtration results for SHP1-C453S. 1C: Turbidity changes of SHP1-WT and SHP1-C453S proteins. 1D: SHP1-WT, SHP1-C453S, and SHP1-E76A proteins purified in vitro (diluted with ddH2O).

**Figure 2.**
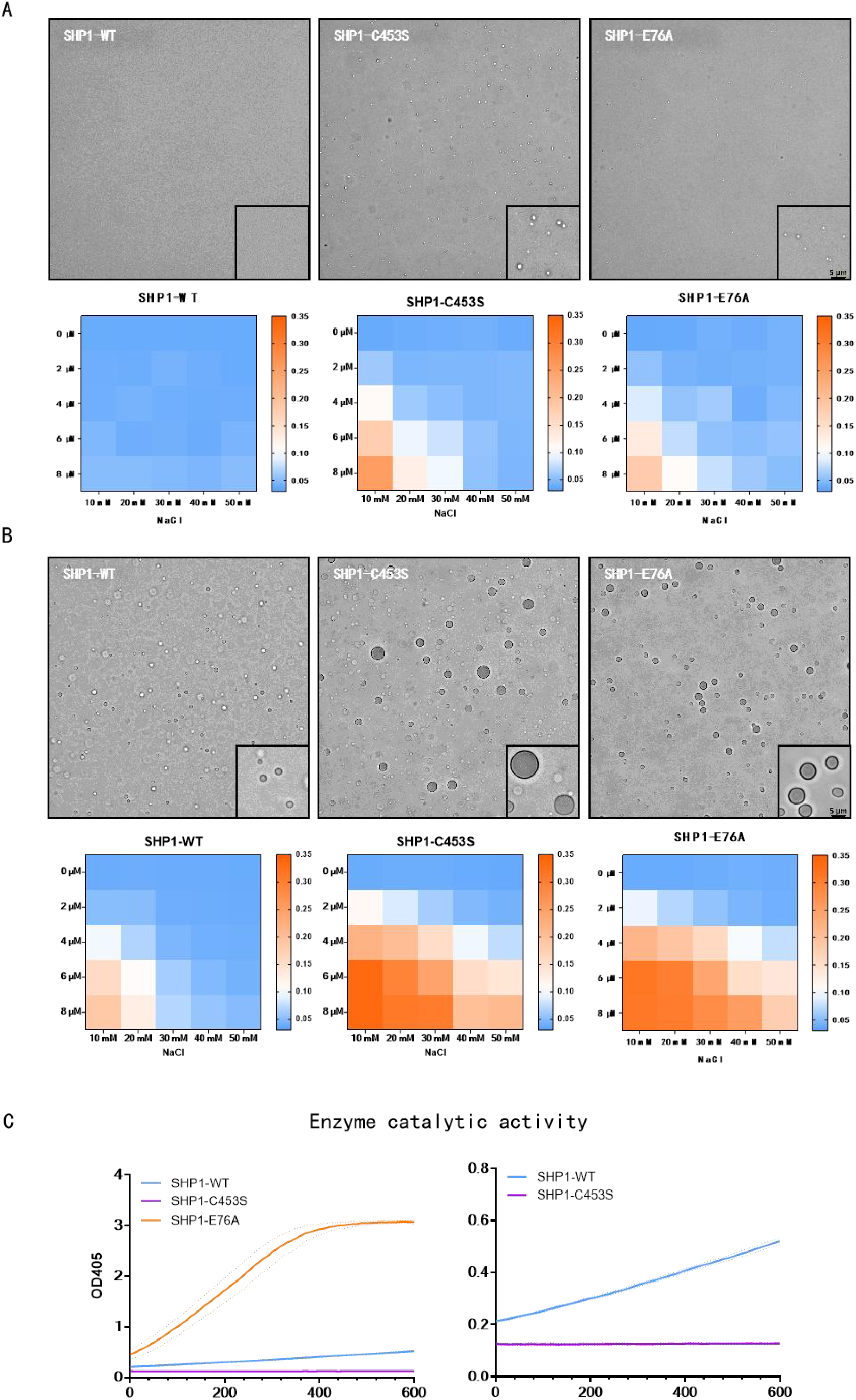
Phase Separation and Enzymatic Activity Testing of Three Proteins: SHP1-WT, SHP1-E76A, and SHP1-C453S. 2A: Representative images of droplet formation by SHP1-WT, SHP1-E76A, and SHP1-C453S proteins, bivariate plots showing the LLPS capability of three proteins at different protein and NaCl concentrations. 2B: Representative images of droplet formation by SHP1-WT, SHP1-E76A, and SHP1-C453S proteins, bivariate plots showing the LLPS capability of three proteins at different protein and NaCl concentrations (in the presence of PEG20000). 2C: Phosphatase catalytic activity of SHP1-WT, SHP1-E76A, and SHP1-C453S proteins by p-nitrophenyl phosphate as substrate.

## Discussion

The C453S mutation in SHP1 may result in a conformational change from a closed to an open state, potentially accounting for the enhanced phase separation capability observed in the SHP1-C453S mutant. Firstly, it has been reported that the inactive C459S mutation in SHP2, a homolog of SHP1, leads to a shift from a closed to an open conformation with greater magnitude compared to other known SHP2 mutants including E76K^[9]^. Secondly, previous studies on phase separation of SHP1/SHP2 have demonstrated that their LLPS activity are positively associated with the extent of openness in their conformation^[8,10]^. Lastly and importantly, structural analysis revealed that SHP1-C453S showed an open conformation rather than the closed conformation observed in wild-type SHP1^[11,12]^. However, the authors were not aware of the potential conformational change induced by C-to-S mutation.

The C-to-S mutation requires particular attention. The structures of cysteine and serine residues exhibit remarkable similarity, with the only difference being a single congener element (-SH changing to -OH). Consequently, in circumstances where it is imperative to disrupt the activity of cysteine without affecting the protein structure, replacing it with serine (S) has traditionally been considered a viable option. However, our work and several other studies ^[9,11,12]^ have suggested that C-to-S mutation in phosphatase could result in a significant structural alterations and impacting its capability of LLPS.

Therefore, a more appropriate mutant is required as an inactive control to investigate SHP1 catalytical activity. The impact of the C453S mutation on LLPS and conformation may potentially result in additional functional changes, thereby influencing subsequent experimental outcomes. It is recommended to identify a novel mutant that produces minimal alterations in structure and LLPS activity while completely abolishing catalytical activity.

## References

[1] Matthews R J, Bowne D B, Flores E, et al. Characterization of Hematopoietic Intracellular Protein Tyrosine Phosphatases: Description of a Phosphatase Containing an SH2 Domain and Another Enriched in Proline-, Glutamic Acid-, Serine-, and Threonine-Rich Sequences[J]. Molecular and Cellular Biology, 1992, 12(5): 2396–2405.

[2] Yi T L, Cleveland J L, Ihle J N. Protein Tyrosine Phosphatase Containing SH2 Domains: Characterization, Preferential Expression in Hematopoietic Cells, and Localization to Human Chromosome 12p12-P13[J]. Molecular and Cellular Biology, 1992, 12(2): 836–846.

[3] Tsui F W L, Martin A, Wang J, et al. Investigations into the Regulation and Function of the SH2 Domain-Containing Protein-Tyrosine Phosphatase, SHP-1[J]. Immunologic Research, 2006, 35(1–2): 127–136.

[4] Garg M, Wahid M, Khan F. Regulation of Peripheral and Central Immunity: Understanding the Role of Src Homology 2 Domain-Containing Tyrosine Phosphatases, SHP-1 & SHP-2[J]. Immunobiology, 2020, 225(1): 151847.

[5] Abram C L, Lowell C A. Shp1 Function in Myeloid Cells[J]. Journal of Leukocyte Biology, 2017, 102(3): 657–675.

[6] Alberti S, Gladfelter A, Mittag T. Considerations and Challenges in Studying Liquid-Liquid Phase Separation and Biomolecular Condensates[J]. Cell, 2019, 176(3): 419–434.

[7] Banani S F, Lee H O, Hyman A A, et al. Biomolecular Condensates: Organizers of Cellular Biochemistry[J]. Nature Reviews. Molecular Cell Biology, 2017, 18(5): 285–298.

[8] Zhang Q, Kong W, Zhu T, et al. Phase Separation Ability and Phosphatase Activity of the SHP1-R360E Mutant[J]. Biochemical and Biophysical Research Communications, 2022, 600: 150–155.

[9] Sha F, Kurosawa K, Glasser E, et al. Monobody Inhibitor Selective to the Phosphatase Domain of SHP2 and Its Use as a Probe for Quantifying SHP2 Allosteric Regulation[J]. Journal of Molecular Biology, 2023, 435(8): 168010.

[10] Zhu G, Xie J, Kong W, et al. Phase Separation of Disease-Associated SHP2 Mutants Underlies MAPK Hyperactivation[J]. Cell, 2020, 183(2): 490-502.e18.

[11] Yang J, Liu L, He D, et al. Crystal Structure of Human Protein-Tyrosine Phosphatase SHP-1[J]. Journal of Biological Chemistry, 2003, 278(8): 6516–6520.

[12] Wang W, Liu L, Song X, et al. Crystal Structure of Human Protein Tyrosine Phosphatase SHP-1 in the Open Conformation[J]. Journal of Cellular Biochemistry, 2011, 112(8): 2062–2071.

